# Yorzoi: Predicting RNA-seq coverage from DNA sequence in yeast

**DOI:** 10.1101/2025.09.20.677345

**Authors:** Timon Schneider, Abdul Muntakim Rafi, Cassandra Jensen, Daniella Liao, Yiren Zhao, Carl de Boer, Tom Ellis

## Abstract

Yeast is one of the principal model organisms in synthetic biology and a widely used chassis for testing heterologous DNA sequences. However, designing DNA constructs correctly is currently limited by our incomplete quantitative understanding of how sequences are transcribed and translated, especially non-native sequences. Here, we present *Yorzoi*, a sequence-to-expression model for bakers’ yeast that predicts RNA-seq coverage for a 5 kilobase, multi-gene window of DNA at 10 base pair (bp) resolution. To extend its predictive ability beyond native DNA, *Yorzoi* has been pretrained on a comprehensive dataset not only including native yeast sequences but also human sequences expressed in yeast and structurally rearranged synthetic yeast chromosomes. We demonstrate that our model has learned general rules of yeast transcription by achieving high predictive performance on various downstream tasks. *Yorzoi* is a powerful tool for *in-silico* testing of DNA sequences and directly applicable for sequence design. A web application to use our model is available at yorzoi.eu and the code open source on GitHub.

## Introduction

Synthetic biology aims to design and assemble genetic constructs that endow cells with valuable functions and capabilities. The yeast *Saccharomyces cerevisiae* is one of the primary eukaryotic model organisms and is widely used in many labs and companies for various applications, ranging from producing valuable molecules to detecting heavy metals [1], [2]. In yeast, transcription is greatly shaped by local DNA sequence features: arrays of transcription-factor binding sites and core promoter motifs set RNA Polymerase initiation and transcription start site usage; sequence-encoded nucleosome affinities (e.g., poly(dA:dT) tracts) gate accessibility; and 3′ terminator signals for cleavage and polyadenylation tune termination efficiency and isoform choice.

Recent machine learning models promise to generate DNA sequences for yeast spanning the most important genetic parts including promoters, coding sequences and full transcripts [3], [4], [5]. However, it is often unclear whether generated DNA sequences are expressed as intended, a key constraint for their practical value in the lab. Generative models now let us sample vast numbers of plausible DNA sequences, but because synthesizing and testing constructs remains expensive, we need computational tools to cheaply prioritize the few designs most likely to express reliably in vivo.

Sequence-to-expression models like Enformer, Borzoi, and recently AlphaGenome predict DNA expression across many modalities, like chromatin accessibility, transcription factor binding and transcriptional activity from DNA sequence directly [6], [7], [8]. Yet existing sequence-to-expression models for yeast are limited to sequences spanning up to one gene and often restricted to predict a single value for the full input sequence [9], [10]. To evaluate and engineer model-generated sequences spanning multiple genes and larger constructs, existing models are inadequate. First, they do not account for the effects of adjacent DNA sequences, which, depending on the intergenic distance, can have a significant impact on local gene expression [11]. Second, transcriptional activity of long sequences must be mapped along the sequence to be useful. For example, transcriptional interference can arise in multi-gene constructs when an upstream transcript reads through a weak terminator and occludes a nearby downstream promoter, when closely spaced promoters collide via opposing polymerases, or when a bidirectional promoter drives unexpected antisense transcription [12], [13], [14], [15]. In these instances, transcript coverage profiles are reshaped without necessarily changing a per-sequence scalar readout (e.g., fluorescence or TPM), so only base-pair- and strand-resolved predictions reveal which parts of the construct are actually being transcribed.

Here, we introduce *Yorzoi*, a Borzoi-inspired deep learning model that predicts normalized strand-specific RNA-seq coverage for the yeast *S. cerevisiae*, for a 5 kb DNA sequence spanning up to 3 genes at 10 bp resolution. For the first time, we supplement a sequence-to-expression model for yeast with RNA-seq experiments that include the expression of human DNA in yeast from artificial chromosomes and strains with a structurally rearranged chromosome, broadening the sequence diversity beyond the native sequence space. In addition to coverage, *Yorzoi* achieves substantial zero-shot performance in predicting MPRA readouts for random core promoter libraries and engineered terminators.

## Results

### Modelling RNA-Seq Using a Deep Neural Network

Similar to other sequence-to-expression models, *Yorzoi*, predicts strand-resolved RNA-seq coverage profiles in the yeast *S. cerevisiae* as a function of DNA for multiple bulk RNA-seq experiments in parallel. It is a deep neural network whose architecture is adapted from Borzoi and Flashzoi (Figure 1 c) [7], [16]. In short, *Yorzoi* consists of convolutional layers that downsample the input sequence, followed by transformer blocks performing bidirectional attention. This captures local motifs (e.g transcription factor binding sites) as well as longer range effects such as promoter-gene context and 5′UTR and 3′UTR interactions. We optimized *Yorzoi* with a shape- and magnitude-decomposed bin-wise Poisson negative log-likelihood, training for 250 epochs with a batch size of 30 on a single NVIDIA RTXA6000 GPU and selecting the best checkpoint by highest log-likelihood on the test-set.

**Figure 1:**
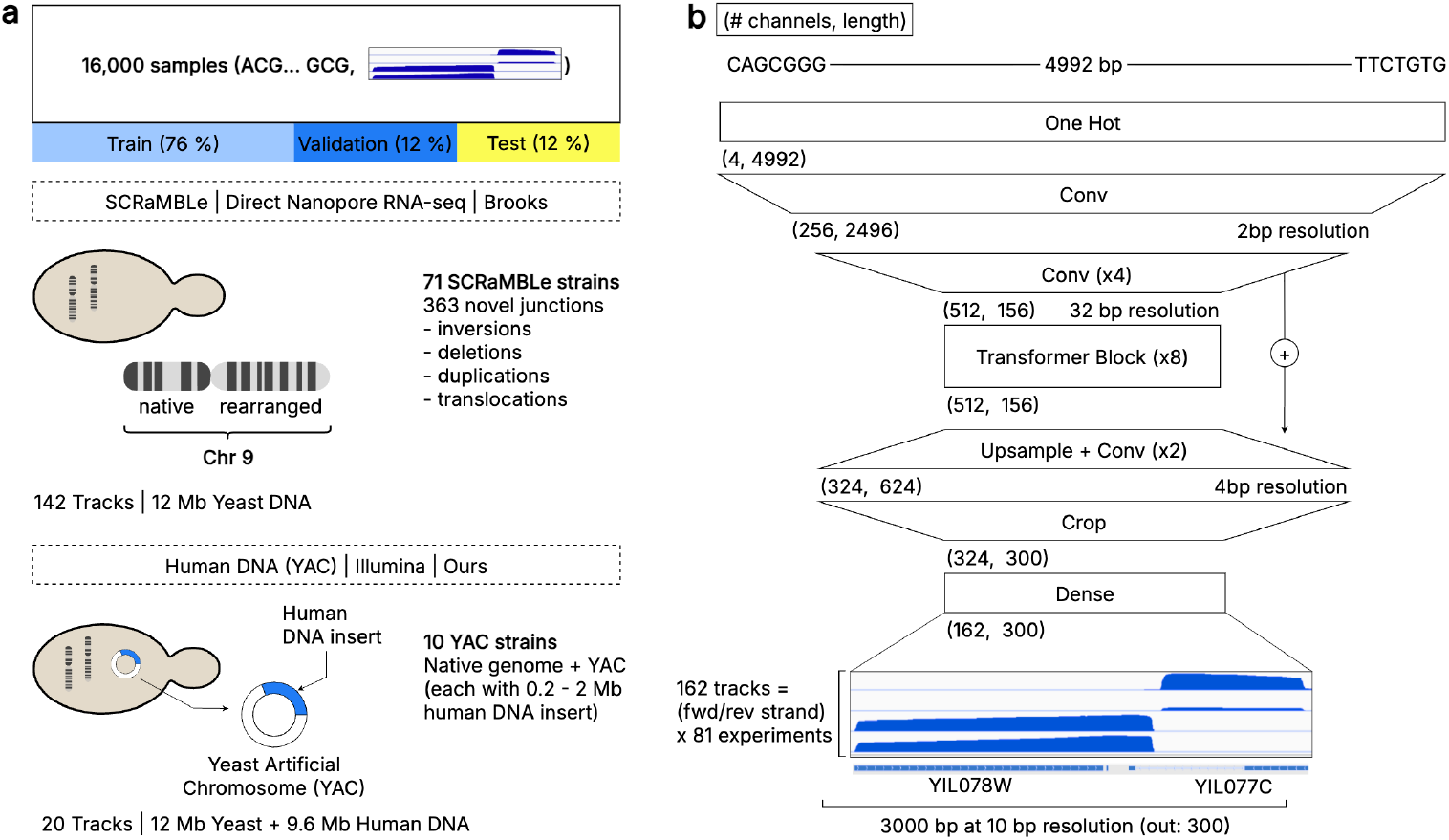
**a** *Yorzoi* predicts RNA-seq profiles from 81 independent RNA-seq datasets. This includes 71 datasets from yeast strains with a SCRaMBLed chromosome 9 (Brooks et al.) and 10 newly generated datasets from strains each carrying a yeast artificial chromosome (YAC) harboring a distinct 200 kb - 2 Mb human DNA insert [11]. **b** *Yorzoi’s* model architecture adapted from Borzoi, predicting the central 3 kb coverage from a 5 kb input DNA sequence across 162 RNA-seq tracks (forward and reverse strand of 81 experiments predicted separately).

*Yorzoi*’s short sequence input window of 5 kb, reduced from Borzoi’s 524 kb, caters to the stark size difference between the yeast and human genome: *S. cerevisiae*’s genome is roughly 500 times shorter than the combined human and mouse genomes that provide Borzoi’s training corpus. Splitting the yeast genome at Borzoi’s input length would yield prohibitively few samples for training. In addition, long range effects (>10 kb), such as enhancers are less relevant in yeast [6], [17]. As spatially proximal (<1 kb) genes influence one another, single-gene windows cannot always explain a gene’s transcriptional variance. We therefore use wider sequence inputs spanning multiple genes [11].

To train our model, we collected sequence-coverage profile pairs from two sources. First, 71 long-read direct RNA-sequencing experiments from *Brooks et al*., which sequenced native and scrambled synthetic sections of the yeast genome [11]. Second, 10 of our own short-read Illumina RNA-seq experiments across 10 yeast strains, each carrying an artificial chromosome with a large human genome DNA sequence insert. Previous work shows that human DNA sequences, which did not evolve in yeast, are abundantly transcribed in yeast with elevated antisense and overlapping transcription, albeit at less extreme levels [18]. We hypothesized that these data would allow our model to generalize to synthetic, non-native sequences, such as those generated for and used in synthetic biology. In total, we work with ≈ 21 Mb of which 12 Mb come from the native genome and its rearrangement and 9 Mb from human DNA sequences in YACs. Tiled into overlapping 5 kb long sections, we obtain ca. 16-thousand samples.

### *Yorzoi* Predicts RNA-seq Coverage

*Yorzoi* predicts 5 kb RNA-seq coverage profiles with a median Pearson R of 0.69 for hold-out sequences (Figure 2 a, c). The hold-out set consists of 12% of the 5-kb sequence windows, stratified across every native chromosome and the human DNA inserts. For each sample, we compute the Pearson R of the predicted profile with its corresponding true profile, track by track, and summarize the resulting correlations by taking the median to derive a single per-sequence Pearson R. While Pearson correlation describes how well our model captures the true shape of the coverage profile, the relative error of the total counts in a prediction window, log transformed,^3^ assesses how well our model captures the magnitude of transcriptional activity: *Yorzoi* achieves a magnitude error that is distributed with a mean *µ* = −0.2 and standard deviation *σ* = 1 for native yeast DNA sequences and *µ* = 0.11 and *σ* = 0.8 for human DNA sequences. In practice, this means that model predictions are expected to be in the range of −56% to +74% of the true transcriptional magnitude for yeast-like DNA sequences.

**Figure 2:**
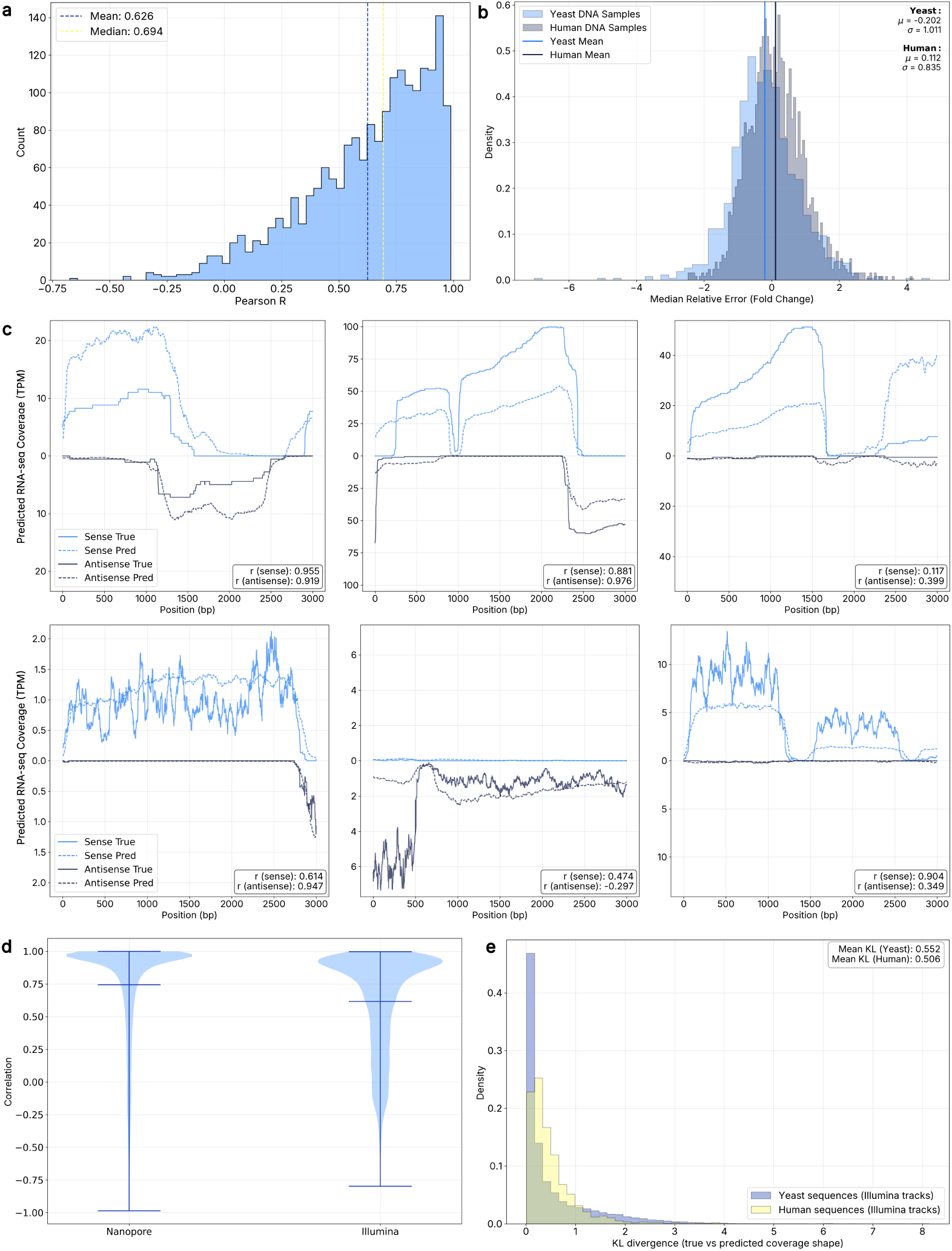
**a** Distribution of Pearson R’s (correlation of predicted with true coverage) aggregated across tracks by median. **b** Track-Median Relative Error (fold change) for yeast and human sequences. **c** Predicted and true coverage values (sense and antisense strand) for six different sequences with Pearson R’s for a selected Nanopore-derived (top row) and Illumina-derived (bottom row) track. **d** Pearson R’s of Illumina and Nanopore tracks point to different biases contained in each sequencing assay. **e** KL divergence (in bits) of count normalised coverage profiles for Illumina-derived sequences partitioned into human and yeast sequences showing yeast DNA sequences to be predicted worse.

**Figure 3:**
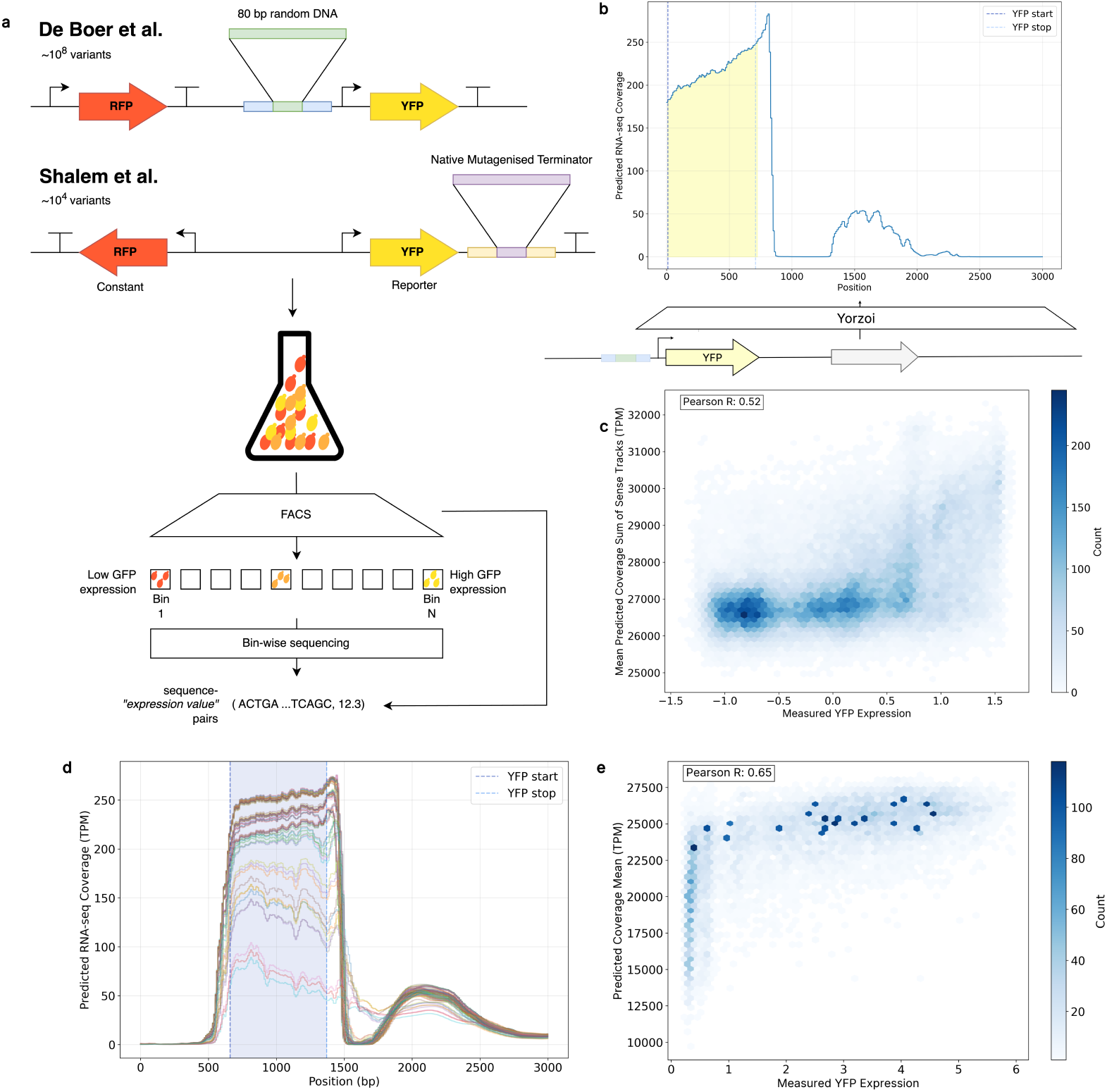
**a** We evaluate *Yorzoi* on two MPRAs, from *de Boer et al*. and *Shalem et al*.. Both generate pairs of sequence and GFP expression for promoter and terminator variants, respectively. While the *de Boer et al*. MPRA works with ~10^8^ random core promoter variants, the *Shalem et al*. measures the expression for ~10^4^ terminator variants [19], [22]. Both assays employ a similar technique of first creating a combinatorial library, which is then FAC-sorted by YFP expression. Bins are subsequently sequenced to map the sequence to expression. **b** For each sequence in the promoter test set, we predict the RNA-seq coverage for the forward strand using *Yorzoi* and sum coverage values for the coding region as a proxy for YFP expression. **c** Predicted coverage sums (Y-axis) are correlated with measured protein expression levels (X-axis), showing a Pearson R of 0.522. **d** *Yorzoi* predicts varying levels of mRNA coverage for terminator variants from *Shalem et al*. [22]. Each line is a different terminator variant and coverage over the blue-shaded region is summed and compared to measured YFP expression levels. Some of the terminator variants show readthrough into the adjacent gene reducing YFP expression. **e** Predicted Coverage sum (averaged across tracks) vs. YFP expression (Pearson R=0.65)

Coverage tracks were derived using two different sequencing methods (Illumina and Nanopore sequencing) that are predicted with varying model performance: Long-read Nanopore data is predicted more accurately (mean *R* = 0.73 on native yeast sequences) than Illumina short-read data (mean *R* = 0.61), presumably because of the experimental bias of Illumina coverage profiles, leading to a more jagged appearance, that is captured less well by *Yorzoi* (Figure 2 d).

Transcriptional landscapes differ markedly between native yeast loci and human inserts: 99% of bases in human fragments in yeast are transcribed - at basal levels and 29% bidirectionally - whereas only an estimated 75% of positions in native yeast DNA show transcription, albeit with more extreme expression levels (high and low) and fewer overlapping transcripts [18]. Thus, we asked whether *Yorzoi* performs equally well on human and yeast DNA sequences. Measured by relative error (*σ*_human_ = 0.8; *σ*_yeast_ = 1) and KL divergence of the count normalized profiles (mean human: 0.5 bits; mean yeast: 0.55 bits), we conclude that differences are present but minimal (Figure 2 b, e).

### *Yorzoi* demonstrates zero-shot performance on promoter and terminator sequence MPRAs

Creating and optimizing promoter sequences is a frequent concern for many applications in synthetic biology. To assess *Yorzoi’s* utility in this domain, we evaluated our model on a promoter-MPRA dataset, which reports a fluorescence readout for millions of unique 80-bp core-promoter sequences [19], [20]. We reasoned that changes in yellow fluorescent protein (YFP) transcript abundance should account for most of the variance in fluorescence levels which would thus provide an appropriate evaluation task for *Yorzoi*. Here, we predict the strand-specific RNA-seq coverage for each of the 70-thousand sequences in the test set. Next, the coverage values in the coding sequence interval (from start to stop codon) are summed to obtain one proxy value of YFP expression per track and per sequence which are then averaged over all tracks to obtain one proxy value per sequence. This proxy value is then correlated with the measured YFP expression. As a result, *Yorzoi’s* predictions explain some part of the expression variance (Pearson R = 0.52) despite the fact that the model was never trained on MPRA measurements or sequences contained in the MPRA dataset (zero-shot setting).

In addition to promoters, synthetic biologists also frequently make use of selected terminator parts from modular cloning toolkits such as the MoClo Yeast Toolkit [21]. Any attempt at scalably expanding the sequence space of possible terminators has to rest on a predictive understanding of sequence determinants of transcription termination. Similar to our promoter MPRA evaluation, we test *Yorzoi* on the 13 k-member 3′-end library dataset of *Shalem et al*., in which each variant terminator was cloned immediately downstream of a GAL1/10-driven YFP and its expression quantified by flow-cytometry for fluorescence measurement [22]. For every construct we padded the full insert to 5 kb and predicted the RNA-seq coverage across the entire region. We summed the coding sequence coverage to obtain one predicted abundance value per variant, treating this as *Yorzoi*’s proxy for steady-state mRNA level. Our predicted proxy correlates with expression levels by Pearson R = 0.65 which indicates that our model has learned predictive rules of transcription termination.

### Capturing the effects of genetic neighborhoods on transcription

*Brooks et al*. find that transcriptional activity changes when a gene is rearranged into a novel genetic context [11]. Across 71 strains, they produce non-native junctions on the right arm of chromosome 9 including swapped 3′ UTRs, gene inversions, translocations and duplications. One example, where a rearrangement changes gene expression is *MND2′s*: here, the RNA-seq coverage over the coding sequence region increases by 2.1x in the rearranged strain ℱ*S606* compared to ℱ*S94*, the strain carrying recombination sites but without induced scrambling.

We evaluated whether *Yorzoi* can predict the effect of such rearrangements on genes held-out from training. Specifically, we compare the actual change in RNA-seq coverage (summed over the coding sequence) for a rearranged gene with the predicted change (Figure 4) and find that the predicted and actual change are correlated (Pearson R=0.33, Spearman *ρ* = 0.32). However, the balanced accuracy of determining whether a change in the genetic neighborhood increases or decreases RNA-seq coverage is 0.62 (random=0.5) pointing to significant challenges of predicting gene expression changes in these scenarios correctly. In summary, *Yorzoi* only partially captures the transcriptional logic of larger genetic rearrangements, and we deem solving this to be an important future direction of this work.

**Figure 4:**
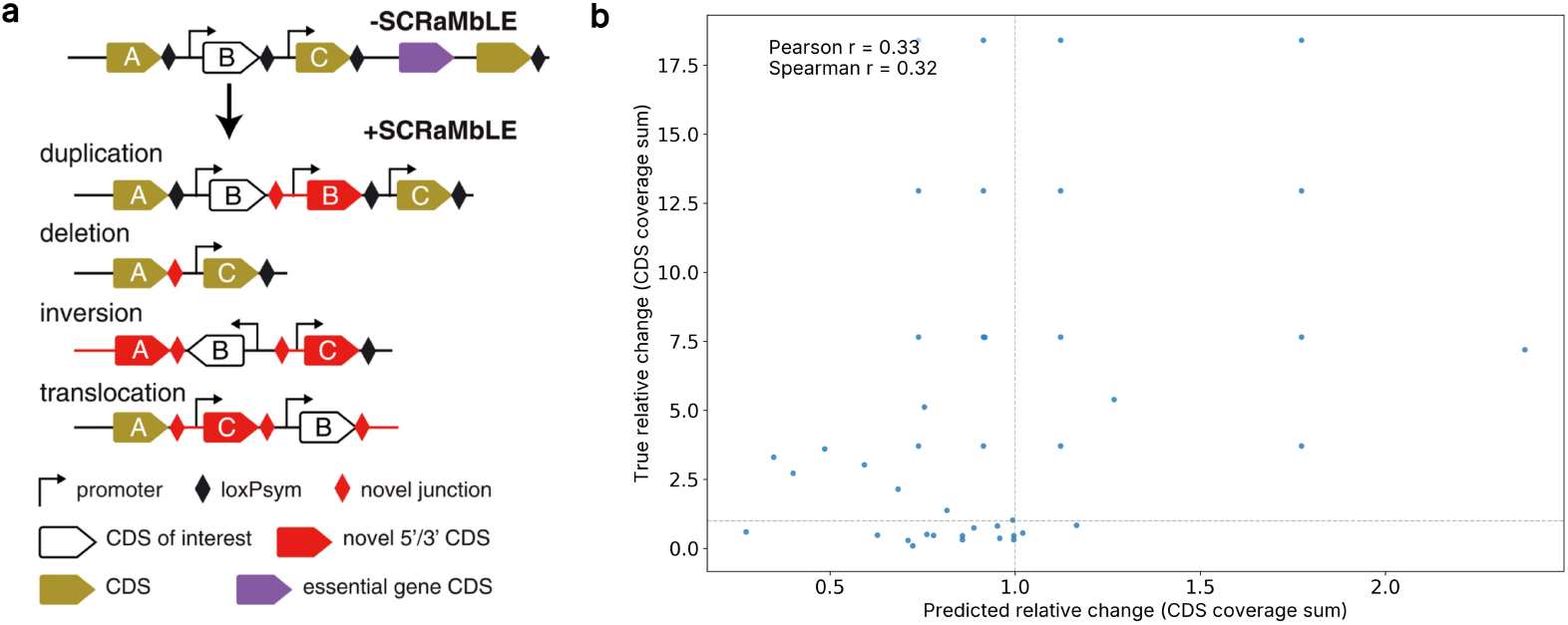
**a** Figure from [11]: example illustration of SCRaMbLE rearranging the genetic neighbourhood of a particular genetic region. The top row of the cartoon shows an example configuration in the unarranged SCRAMBLE strain ℱ*S94* which contains loxPsym recombination sites after every non-essential coding sequence on the right arm of chromosome 9. Upon induction, recombination produces duplications (with novel 3′UTRs), deletions, inversions and translocations. **b** Predicted vs. true relative change in RNA-seq coverage summed over the coding sequence region for all genes rearranged into a novel context, i.e. relative_change 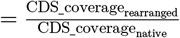. For example, a relative change of 0 refers to a change where the rearranged coding sequence is not transcribed at all. The largest gain in transcriptional activity is of *VLD1* which is transcribed ~17.5 fold more than its unarranged counterpart. Dotted lines indicate a relative change of 1 (i.e. no change in transcriptional activity). Each of the 41 dots refers to one pair of predicted and true relative change for a single coding sequence (18 unique coding sequences across 5 strains).

**Figure 5:**
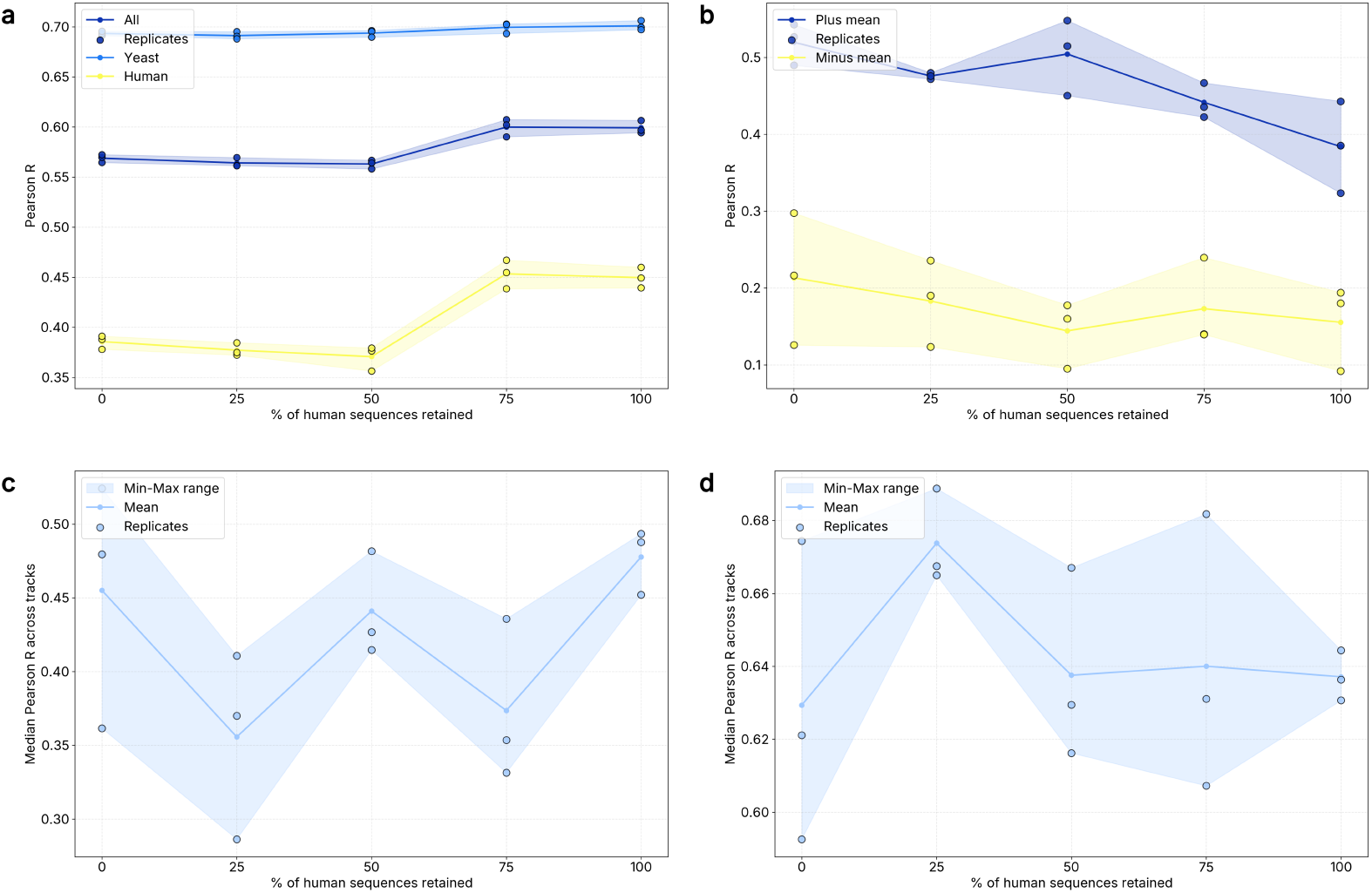
Ablation results from three replicates across five ablation levels where 0% indicates training on yeast sequences only and 100% training on yeast and all available human DNA samples (total: 6409 human sequences). The performance of the individual replicate is shown as circles. The blue mean line depicts the mean of all replicates. **a** Pearson Correlation of true and predicted coverage profiles improves for human DNA sequences with additional human DNA sequences while predictions on native sequences remain largely unchanged. **b** Pearson correlation of samples from *Brooks et al* containing novel junctions, introduced by SCRaMbLE, decreases when the more human DNA sequences are added. **c** Correlation of measured YFP expression levels and predicted coverage sum for the *de Boer et al*. MPRA. Correlation non-monotically changes but improves in mean with increasing number of human DNA sequences. [19] **d** Correlation of measured YFP expression levels and predicted coverage sum for the *Shalem et al*. terminator MPRA. Here, correlation moves in a narrow range from 0.63 to 0.68 and does not improve monotonically. [22].

### Influence of Human DNA Sequences on Model Performance

As part of our study we ran our own RNA-seq experiments profiling 10 yeast strains, each carrying human DNA inserts. Unlike native DNA, human DNA has not evolved in yeast and so is ‘naive’ and shows markedly different expression patterns compared to native DNA. Thus, we hypothesized that supplementing the existing native sequence dataset with data from human sequences should lead to a more general model capable of predicting across a wider sequence context. To test this, we ran an ablation study with three replicates, adding successively more human DNA sequence samples into the training (0%, 25%, 50%, 75% and 100% of human DNA samples). For our training task of predicting coverage profile, performance improves correlations of true and predicted coverage for both sequence origins, yeast and human, but has a stronger positive effect for predicting human RNA-seq coverage. However, our results do not show that adding human DNA sequences into the training generally improves performance on downstream tasks such as predicting the effect of promoter or terminator variants on gene expression. Especially the deteriorating effect of increasing the number of human sequence samples on predicting the coverage of genes rearranged into novel contexts remains an open question.

### Application of *Yorzoi* for Synthetic Biology Applications

The best way to use *Yorzoi* without any coding is through our web interface at Yorzoi.eu. To illustrate how we think it could be used in synthetic biology, we have tested it for three uses described below; (i) predicting the effect of using different modular promoters and checking the effect of altering gene order and orientation, (ii) evaluating different codon optimizers and (iii) using it for sequence design [21].

#### Use Case i

Gene order can shape gene expression by placing transcription start sites in different upstream contexts, yet synthetic biologists seldom know which arrangement will work best. Using *Yorzoi*, we compared alternative orders for a five-gene pathway that turns tryptophan into violacein. Figure 6 a–b show that surrounding context matters: vioD coverage drops when it is placed divergently from vioA but recovers when it leads the construct. Digital evaluation of gene order, promoter strength, and gene orientations like this can flag design pitfalls before in-vivo testing.

**Figure 6:**
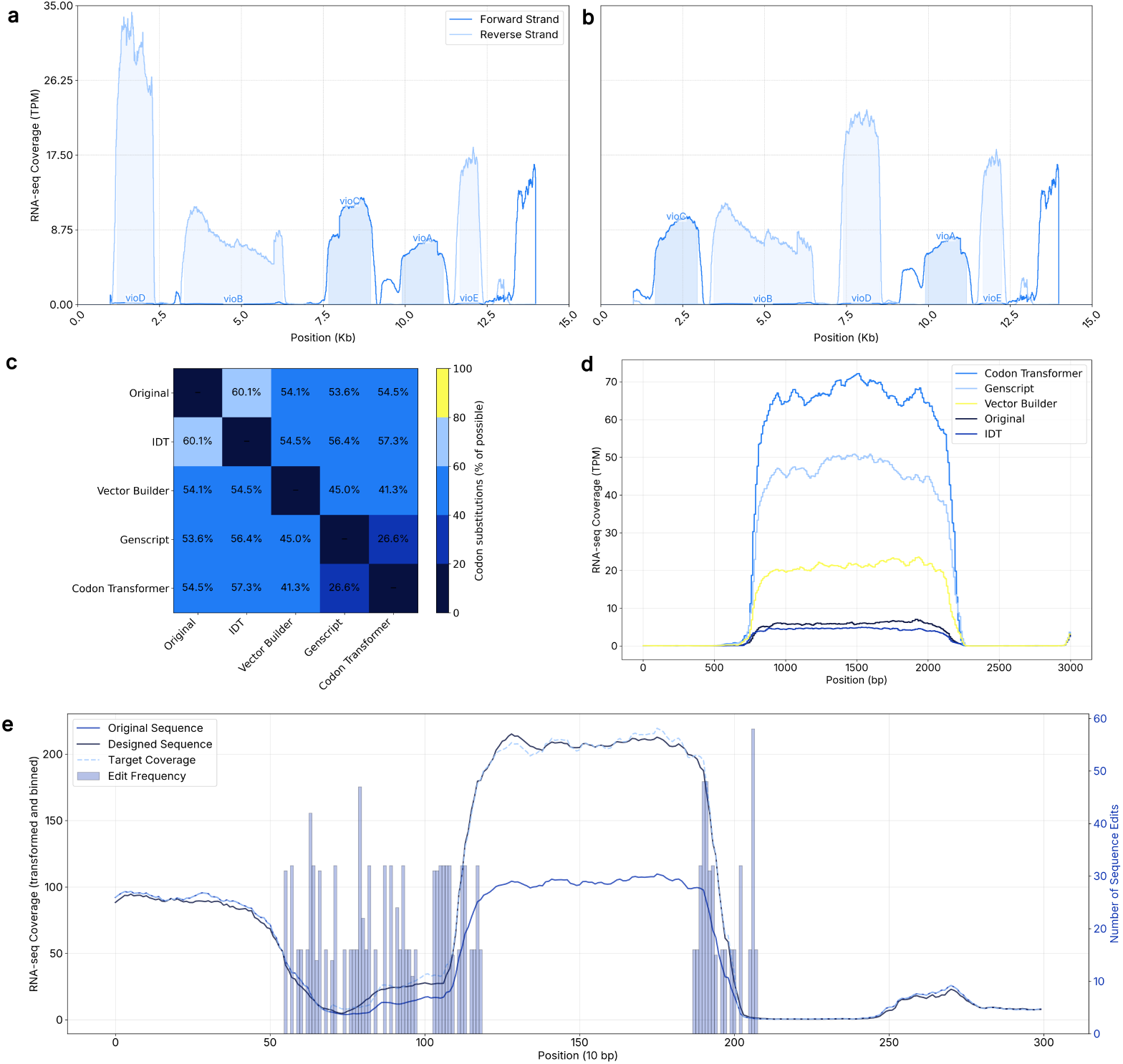
**a** and **b** showing two different constructs of a 5-gene violacein cluster (vioA to vioE) where only the order of the genes was changed which reveals substantial differences in predicted expression levels for vioD when put in divergent orientation to vioA. Shaded regions delineate the coding sequence region of each gene. **c** Matrix showing the percent of codons substituted for synonymous ones by different codon optimizers. **d** RNA-seq coverage for five different vioC gene constructs where only the coding sequence was varied using synonymous codons substitutions suggested by four codon optimizers. Compared to the original sequence, expression is expected to vary by 15-fold. The legend is ordered in the same order as the coverage levels. **e** To design a new sequence, we constrained Ledidi to only make edits in the direct upstream and downstream regions of the GFP coding sequence. The histogram bars show the number of edits made for a single designed sequence to achieve the target coverage for one of the designed sequences. The three coverage lines show the predicted RNA-seq coverage of the original DNA sequence containing the GFP coding region, the target coverage and the predicted coverage for the Ledidi-edited sequence closely matching the target coverage.

**Figure 7:**
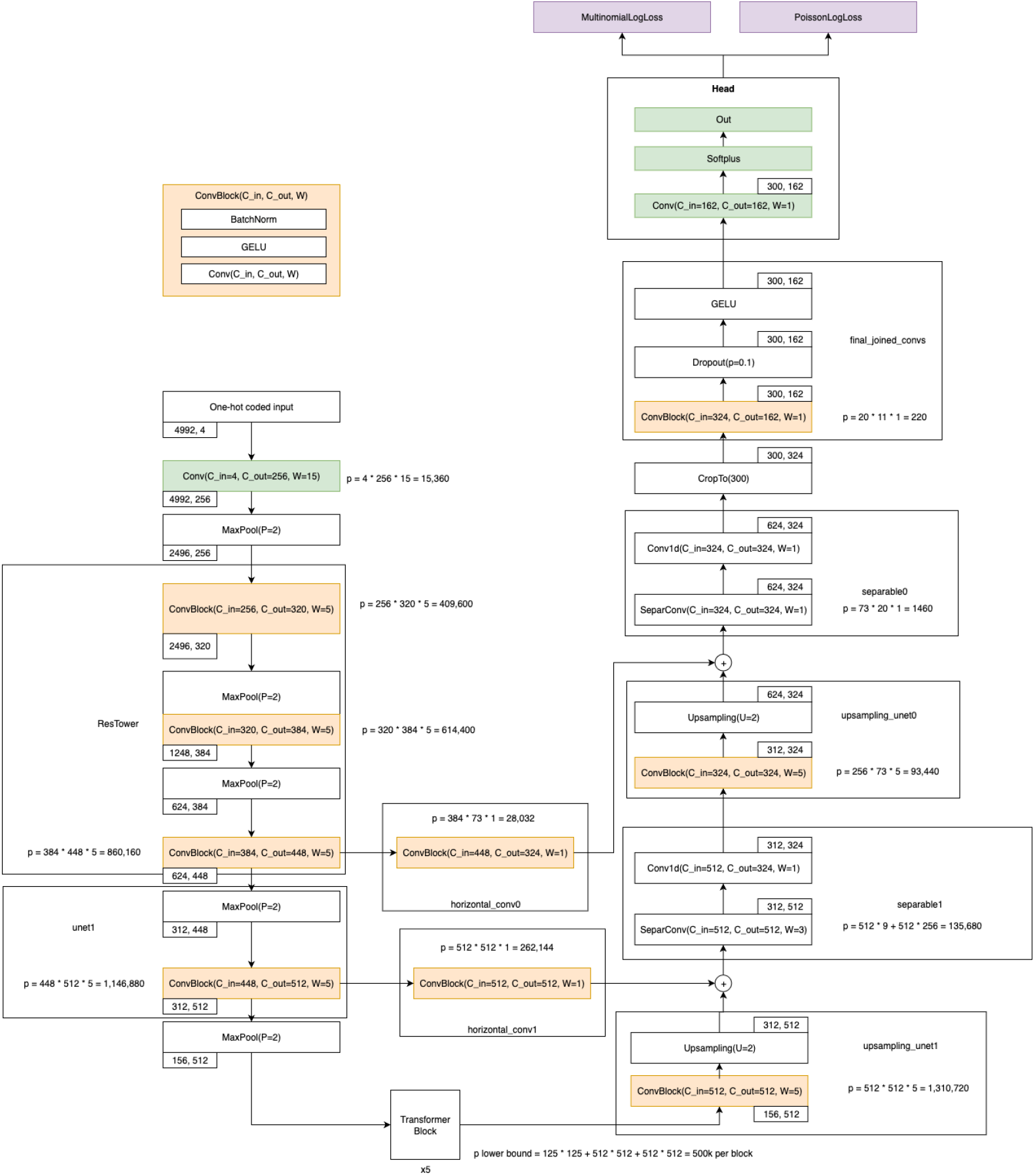
*Yorzoi* full model architecture for ~ 5 kb input window and 162 tracks (81 RNA-seq experiments).

#### Use Case ii

Codon optimization is one of the most frequently used computational tools in synthetic biology but the effect of optimizations on transcription mostly remains unclear. Yet, changes to the codon sequence can have unintended effects on chromatin organization or introduce premature polyadenylation signals [23]. With *Yorzoi* researchers can predict the effect of codon optimization on transcript levels. To illustrate this, we tested four different codon optimization algorithms available online, by using *Yorzoi* to predict coverage of different designs where only the coding sequence of a gene was varied. As an example, we show that commonly used codon optimizers generate diverse coding sequences with up to 60% of codons being substituted for synonymous ones and reveal that these different designs are predicted to vary transcript levels by ~15x (Figure 6 c,d).

#### Use Case iii

*Ledidi* is a framework that allows users to employ sequence-to-function models, like *Yorzoi*, to design sequence edits that change an initial sequence to one with user-specified coverage levels [24]. With *Yorzoi* researchers can now use Ledidi to design sequences for yeast that have a target RNA-seq expression level. As an example, we sought for edits in the non-coding regions upstream and downstream of a GFP-encoding protein that would increase GFP transcript levels by two-fold. Figure 6 e shows the resulting sequence edits in the regulatory regions adjacent to the coding sequence. Figure 6 f shows how the suggested sequence edits cause the predicted expression level to double and match the intended target expression.

Looking forward, we hope to enhance *Yorzoi* with features relevant to synthetic biologists. For example, predicting and designing promoters that are strictly off in general conditions and on in induction conditions would be very useful, but would likely require adding more experimental results to the current training sets.

## Discussion

Here we introduce *Yorzoi*, the first deep learning model designed to predict RNA-seq coverage in the eukaryotic model organism *S. cerevisiae* from DNA sequence at a near-base pair level resolution spanning multiple genes. Across 81 RNA-seq experiments, including human DNA sequences as well as synthetic SCRaMbLE yeast chromosome strains *Yorzoi* makes relatively accurate predictions on hold-out test sequences. Training on non-native DNA sequences, notably human genomic inserts in yeast artificial chromosomes and structurally rearranged synthetic genome sections, extends *Yorzoi* predictions beyond the native sequence space. We also demonstrate that *Yorzoi* generalizes to novel assays such as promoter and terminator MPRAs without additional finetuning on MPRA data.

Key challenges remain in modelling structural rearrangements, such as those produced by SCRaMbLE of synthetic chromosome arm 9, and also in modelling data from Illumina experiments. As seen in our ablation study, the contribution of human DNA transcription data to improved performance on downstream tasks is only sometimes marginally positive, which runs against our initial expectations that for this given model architecture, more, diverse, transcription initiation and termination sequences would lead to better generalisation to other unseen sequences such as those tested in the MPRA. We currently attribute this to shortcomings in either the chosen model architecture or the training method.

Future research could systematically evaluate a range of new architectures and training methods to achieve more consistent improvements across tasks, including those involving heterologous DNA sequences. To increase utility for researchers, datasets representing a more diverse range of conditions - such as varying carbon sources and growth conditions - should be assembled and used as training targets. The near-completion of the Sc2.0 synthetic yeast genome project may provide a valuable source of data for addressing current limitations in modelling RNA-seq coverage of structural genomic rearrangements. Diverging from standard coverage-prediction approaches, future efforts might instead model read data directly to preserve transcriptional logic that coverage profiles can obscure - for example, in overlapping transcription frequently observed at novel junctions [11].

In summary, we have built the first AI model predicting RNA-seq coverage profiles from DNA sequence for yeast and show its zero-shot performance on MPRAs, enabling *in-silico* design and screening of non-native DNA sequences.

## Methods

### Problem Formulation

#### On RNA-seq

For a given genomic sequence of an organism, in our case *S. cerevisiae*, bulk RNA-seq experiments detect a subset of mRNA molecules synthesized by a bulk of approximately genetically identical cells. After isolating RNA, the molecules are sequenced resulting in millions of reads. We map each read back onto the reference yeast genome (read alignment) and record the overlapping reads at each genomic sequence position for both strands separately which gives us two strand-specific *coverage profiles* (Figure 1 a). As such, coverage profiles resolve transcriptional activity spatially and indicate its magnitude.

#### Mathematical Formulation

In line with previous work, we frame predicting RNA-seq assays as a genomic track prediction problem (Fig 1 b):

Consider the i-th sample input, a DNA sequence *S*^(*i*)^ ∈ {*A, C, G, T* }^*L*^ (e.g. AGTGACGGA…) where *L* is the sequence length and corresponding 2*M* normalized RNA-seq coverage tracks from *M* RNA-seq experiments where each experiment contributes two tracks, one for the positive strand and the other for the negative strand coverage.

*T*^(*i*)^ ∈ ℝ^2*M*×*K*^ then describes a matrix of stacked coverage profiles for sequence *i* where *K* is the length of the coverage profile when aggregated at resolution 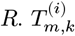 represents the measured normalized coverage value for track *m* at the pseudo-position *k* with

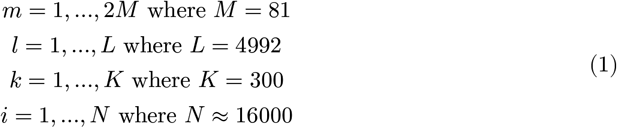

Typically, each character in a DNA sequence corresponds to one coverage value for each track. Here, we restrict the tracks to the central *L* − *R* ∗ *K* positions at a *resolution* of *R* = 10. I.e. the original track of length *L* is cropped on both sides by 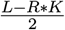 and remaining coverage values are tiled into *K* blocks, *W*_*k*_, of length *R* starting at sequence position *l*_0_:

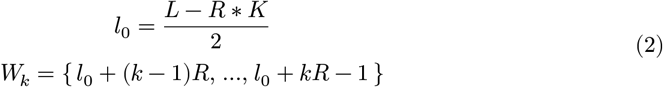

The coverage values within a block are summed to one scalar which we term the normalized read count value on track *m* at *pseudo-position k*. Let *t*_m,l_ ∈ ℝ^+^ be the read count value for track *m* at base *l*:

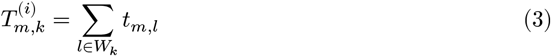

Then, our neural network *f*_θ_ outputs a non-negative tensor:

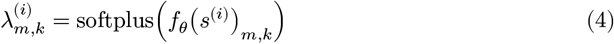

We then additionally assume for the data generating process that

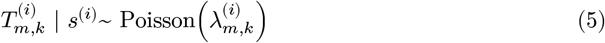

I.e. the normalized RNA-seq coverage values are conditionally independent, given a sequence *S*^(*i*)^

### Data

The data, paired DNA sequence and RNA-seq experiments, comes from two sources, one being *Brooks et al*. [11] and the other being our own RNA-seq experiments. *Brooks et al*. contains 70 strains which differ by their chromosome 9 right arm sequence but are else identical with the native R64-1-1 *S. cerevisiae* reference sequence. The sequences of the right chromosome 9 arm have been experimentally rearranged by inducing recombination at loxPsym sites, placed downstream of the coding sequence of every non-essential gene. The strains also contain the native chromosome 9 left arm sequence. RNA-sequencing was done using direct Nanopore sequencing which produces long reads covering full transcripts. For each strain, the paper authors provided us with the genomic and read files. *bedtools* is used to produce strand-specific coverage files which are normalized by dividing the raw coverage by the total read count and multiplying the resulting relative count by 10^6^. For model training, coverage values are first binned by summing 10 consecutive coverage values and then, similar to the squashing used for training *Borzoi* in [7], normalized RNA-seq coverage values are transformed using the following function:

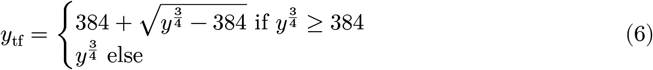

While training is done on binned and transformed values, for all evaluations and downstream tasks predicted are first inverse transformed and then unbinned. The inverse transformation is:

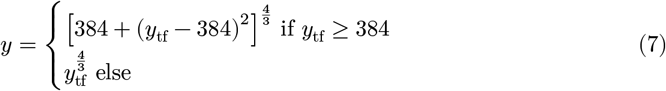

The implementation of both functions can be found on Github (transform, inverse). We additionally conducted our own stranded Illumina RNA-seq experiments on 10 strains which each carry a human DNA insert in a yeast artificial chromosome. All were grown in synthetic defined media lacking uracil. All details of the human DNA inserts can be found in the Supplementary Files.

#### Train-Test Splitting

Samples are derived from the genomic sequence by splitting the genome into non-overlapping sections of length 3 kb. This yields ~16,000 samples. One kb is added upstream and downstream to extend the sequence length to 5 kb. The start site for all 3 kb regions is algorithmically determined by scanning for changes in coverage which means that transcriptional activity tends to start at the left boundary of the 3 kb region. This poses a potential risk for position bias where some part of the prediction is determined by the position of the sequence within the window instead of the sequence features. We test this with a label-shift sensitivity analysis. Model the normalized prediction as

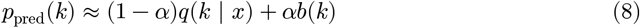

Keeping *p*_pred_ fixed, we circularly shift the labels by Δ bins within the 3 kb window and recompute the multinomial (shape) loss:

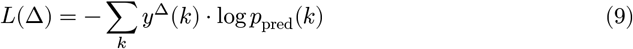

with

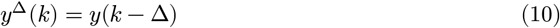

A position template would move the minimum to Δ ≠ 0; in our data both halves minimize at Δ = 0 (Suppl. Fig. S1), indicating negligible position-offset bias.

We divide the samples into two sets, *train* (76%) and *val-test* (24%), by selecting one window on each chromosome (including YACs) which minimizes homology between train and rest. The *val-test* fold is then randomly subdivided into validation and test set. We find the window which minimizes homology by computing ~128 million pairwise homologies (BLAT score) between samples (5 kb sequence) to build a homology graph where nodes are samples and edges indicate homology between two samples with a weight. For each chromosome consecutively, we then slide a window across its genomic sequence and sum the outgoing edges from samples within the window to any other node not in the window (across all chromosomes). This produces one *val-test* window per chromosome which minimizes homology across all sequence samples. This is done to prevent performance overestimation through train-test leakage where for example, similar sequences on two different chromosomes are seen in train and test set.

### Model Architecture

We adapt the Flashzoi architecture for *Yorzoi* [16]. *Flashzoi* is a modified architecture from *Borzoi* that replaces standard cross-attention with FlashAttention2 for higher memory-efficiency and the custom positional encoding with rotary positional embedding and implements the resulting architecture in PyTorch [7], [25]. The general architecture consists of four parts: 1) convolutional layers that downsample the sequence length while increasing the number of channels, 2) 8 transformer blocks (sequence input length: 156) performing cross-attention operations for longer range interactions such as neighboring genes, 3) subsequent upsampling layers with incoming skip connections from the downsampling layers and 4) a head that is finetuned to the specific track.

Yeast genes are typically shorter (*S. cerevisiae*: 1020 bp median length; 90th-percentile 2668 bp vs. *Homo Sapiens*: 24 kb median length) and spaced more densely throughout the genome (Median Integenic Distance - *S. Cerevisiae*: 350 bp vs. *Homo Sapiens*: ~10s of kb) [26]. Since neighboring yeast genes influence each others transcription we pick an input window size of 4992 bp that allows us to fit 1-3 genes to strike a balance between accounting for neighborhood effects and keeping the window size small enough to learn from a sufficient number of samples [11]. To avoid cases where transcription outside the sequence window influences coverage values within the window, we crop the predicted coverage window to the central 3 kb.

In comparison to *Borzoi*, we reduce the model size and computational requirements further by removing layers in the downsampling layers resulting in cross-attention operations at a resolution of 32 bp.

### Training

On an NVIDIA RTXA6000 GPU, training took approximately 1.5 hours. Thanks to the small model size, we could increase the batch size during training to 30 samples from the per-GPU batch-size of 1 that *Borzoi* uses. Throughout training we used a cosine annealing schedule with warm restarts which we found to improve performance when combined with AdamW as the optimizer. We applied reverse complement augmentation randomly in 50% of the cases. In practice this means reverse complementing the input DNA sequence and swapping the 5′ to 3′ tracks in the upper half for their corresponding 3′ to 5′ track in the lower half and reversing the coverage values.

#### Loss

Similar to *Borzoi, Yorzoi* is trained using a loss function that decomposes the loss into a magnitude loss which penalizes the difference between the summed counts across the predicted coverage and a shape loss which penalizes the difference in how reads are distributed across bins. This allows for a reweighting of the shape and magnitude term of the loss. In comparison to *Borzoi* we increased the weight of the shape loss. Our choice for the magnitude loss is the Poisson Negative Log Likelihood and Multinomial Negative Log Likelihood as introduced by BPNet [7], [27]. The exact loss function is as follows:

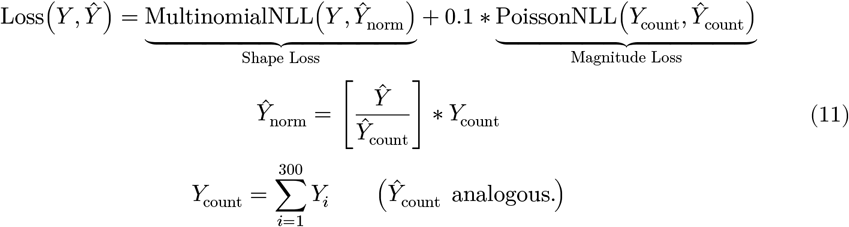

where *Y* ∈ ℝ^300^ is a single true coverage track and *Ŷ* is the predicted one. The full implementation of the loss function can be found on [GitHub](https://github.com/Tom-Ellis-Lab/yorzoi/blob/4f95775d52c695409d4a0dd4638c5c76b60e4e29/yorzoi/loss.py#L5). Its noteworthy that the per-bin Poisson independence assumption is violated in our training data, because individual reads typically span multiple bins, leading to pronounced over-dispersion, characterised by an excess of zero-count bins and heavier-than-Poisson tails. We believe that this did not negatively influence the training process. For sequences with only one corresponding RNA-seq experiment, for example novel junctions from SCRaMbLEd strains or human DNA inserts, we compute a selective loss which only computes the loss on the two available tracks (the two strands). During training, we upweight samples with fewer tracks to have the same loss contribution as samples with more tracks.

### Evaluations

#### Pearson Correlation and Fold-Change Error

For calculating the Pearson correlation in Figure 2, we first unbin and untransform our predictions. Then, we compute the correlation for every track-sample, i.e. we derive one correlation per track for every sample at single base-pair level. Then, we average the correlations over all tracks for a given sample. This reduces the reported performance but represents the performance across sequence more accurately as human DNA sequences are typically only represented on two tracks which would down-weight their influence when simply plotting the histogram over all track-samples. The relative error, reported as fold change, is computed on unbinned and untransformed predictions by averaging the fold change error across the sequence for every track-sample. The fold change error for a true coverage value *Y*_*i*_ and predicted coverage value *Ŷ*_*i*_ at sequence position *i* is simply 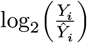 which is equivalent to computing the difference on log transformed data.

#### De Boer et al. Promoter Set

We obtain the test dataset of ~70,000 80 bp promoter sequences and construct sequences by inserting the 80 bp insert into the sequence template obtained from Addgene.org which is the plasmid sequence used for MPRA experiments in [19]. We position the start codon at position 1000 aligning with the start of the 3 kb predictive window of *Yorzoi*. We then predict the coverage for all sequence constructs and sum the coverage of the GFP coding region for each track individually. We average the coverage sums across all forward strand tracks per sample and compute the Pearson R between the averaged coverage sum and the measured YFP expression value.

More precisely, we construct a single input sequence for predicting the coverage as follows:

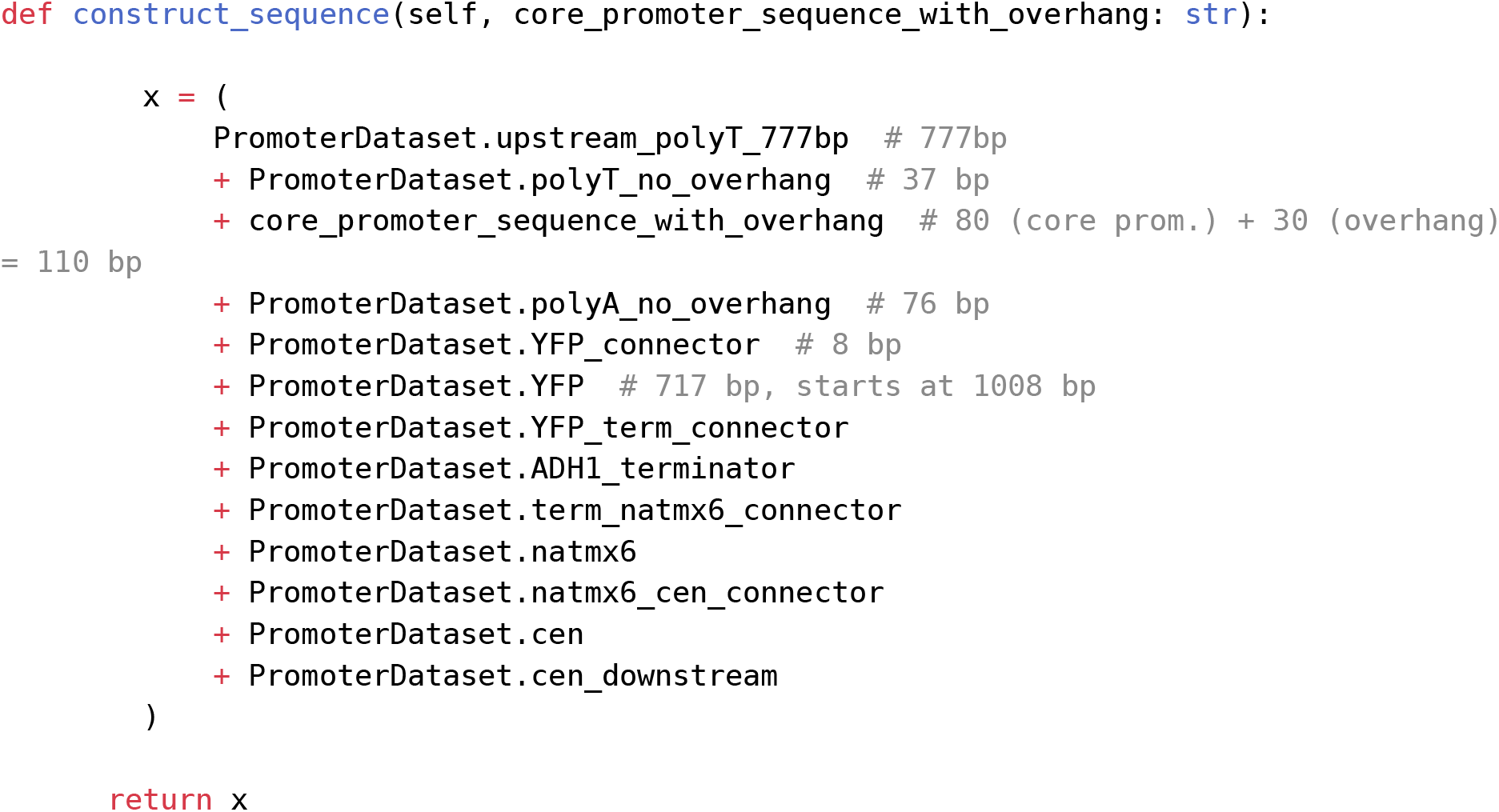

where *x* ∈ *X* is the input sequence for our model, *Yorzoi, f*, to predict the coverage *c, f* : *X* → *C* where *C* = {*c* | *c* ∈ ℝ^162×300^}. We unbin and untransform as described earlier to get *c*^′^ ∈ *C*^162×3000^ and sum over the YFP coding region:

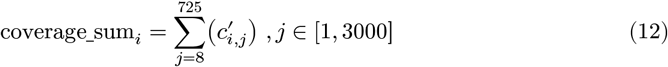

and then average that coverage sum across all sense strand tracks:

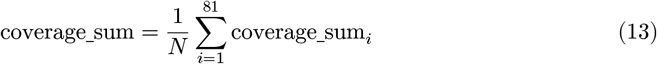

This per-sample coverage sum is then Pearson correlated with the measured YFP expression values.

#### Shalem et al. Terminator Set

From [28], we obtain the data for [22] and perform testing on all sequences. We do this by building a DNA construct consisting of a leading GAL1-10 promoter followed by a YFP coding sequence, the terminator insert and a randomly picked fixed sequence from the coding sequence of CYC1 (as specified in [22]). Then, we use *Yorzoi* to predict RNA-seq coverage for all constructs and sum the coverage across the YFP coding sequence region. We then compare this coverage sum with measured expression levels to compute a correlation for each track. Out of all tracks, we average all to report performance. This is identical to the approach for the *de Boer et al*. MPRA. To demonstrate zero-shot performance, we do not train on the assay data.

### Brooks et al. Novel Junction Set

From the supplementary material of [11], we obtained a table science.abg0162_table_s3.txt containing all determined novel junctions defined as loci where a rearrangement of DNA section could be identified using whole genome sequencing (annotated in the txt file). From this, we identified all genes rearranged into novel context (total 18) across 5 strains yielding a total of 41 samples. For all selected samples, we extracted their coding sequence RNA-sequence coverage of the strand that the gene sits on from the unscrambled strain ℱ*S94* and the respective rearranged strain (one of ℱ*S606*, ℱ*S707*, ℱ*S711*, ℱ*S731*, ℱ*S732*). The coverage is summed to obtain a pair (ℱ*S94 coverage sum, SCRAMBLE strain coverage sum*) of CDS coverage sums per sample. Then, the relative change for each sample is computed as

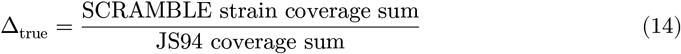

Δ_pred_ is computed by predicting the coverage of all 18 coding sequences (when positioned centrally in prediction window) and predicting the coverage of all coding sequences in their respective rearrangements across the 5 strains. As before the coverage over the coding sequence region is summed and the relative change for a given sample computed as

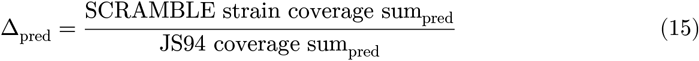

### Human DNA sequence ablation

We trained three model replicates each with a different random seed (torch.manual_seed(x)) for 250 epochs and evaluated the best model as measured by validation loss for each replicate independently. We trained models using 0, 25, 50, 75 and 100 percent of available human DNA sequences (100% = 6409 human DNA sequences) resulting in a total of 15 models.

We evaluated models across four tasks:

1. Pearson corrleation of predicted and true coverage stratified by sequence origin (human or yeast)
2. Pearson correlation of predicted and true coverage of novel junction samples (from SCRAMBLEd chrom 9 right arm)
3. Pearson R of Promoter MPRA performance (as specified in *Methods - De Boer et al. Promoter Set*)
3. Pearson R of Terminator MPRA performance (as specified in *Methods - Shalem et al. Terminator Set*)

### Yeast Artificial Chromosome RNA-seq Experiments

#### Yeast strains and culture conditions

10 YACs with human DNA inserts were ordered from the centre for applied genomics (TCAG) as yeast extract peptone dextrose (YPD) stabs. To select for yeast containing a YAC, yeast cells were streaked onto synthetic defined (SD) plates lacking uracil and containing adenine sulfate (Sunrise Science) and were grown at 30 °C for 2 days. Colony PCR was performed on red colonies to confirm the presence of the human YACs.

#### Yeast YAC RNA-seq experiment

Yeast colonies containing the YACs were grown in liquid SD-URA to an optical density at 600 nm of 0.3–0.6. Total RNA was extracted from liquid yeast cultures using the hot-acid phenol method, described previously75. Genomic DNA was removed from RNA using Turbo DNase following the manufacturer’s instructions (Invitrogen). Extracted RNA was prepared for strand-specific sequencing by the Biomedical Research Centre Sequencing Core at the University of British Columbia using the NEBNext Ultra II Directional RNA Library Prep Kit for Illumina and poly-A+ selection (NEB). Libraries were sequenced on an Illumina Miseq machine using 300 cycles, generating 20 million paired-end reads.

## Weights, Data and Code Availability

*Yorzoi* is accessible for free at yorzoi.eu, no coding or GPUs required.

Code and model weights are available on GitHub and Huggingface:

- Code: https://github.com/Tom-Ellis-Lab/Yorzoi
- Model weights: https://huggingface.co/tom-ellis-lab/Yorzoi

The dataset will be published soon.

## Contributions

DL conducted RNA-seq experiments on YAC-strains. CJ did the bioinformatics work on the YAC strains. AMR consulted and contributed very detailed feedback throughout the process of conceptualising and executing this project. TS conceptualised the project, wrote the code and manuscript. CB, TE and AZ provided feedback and funding for this project.

## Acknowledgements

We want to especially thank Johannes Hingerl for porting Borzoi to Pytorch and making it more easily trainable via *Flashzoi* whose code we adapted for *Yorzoi*, Johannes Linder for early feedback and patiently answering early questions, Maciej Wiatrak and Felix Teufel for detailed feedback on the manuscript, Patrick Kidger for feedback on our repository and Lucas Coppens for his ideas and feedback. We also want to thank the Google Cloud Education Program and Modal Labs for providing compute for model training and data processing. Aaron Brooks kindly provided the data from [11].

## Footnotes

3 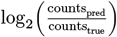

## Bibliography

[1] M. Parapouli, A. Vasileiadis, A.-S. Afendra, and E. Hatziloukas, “Saccharomyces cerevisiae and its industrial applications,” AIMS Microbiology, vol. 6, no. 1, pp. 1–31, Feb. 2020, doi: 10.3934/microbiol.2020001.

[2] E. Wahid et al., “Immobilized Saccharomyces cerevisiae viable cells for electrochemical biosensing of Cu(II),” Scientific Reports, vol. 15, no. 1, p. 2678, Jan. 2025, doi: 10.1038/s41598-025-86702-8.

[3] Z. Li et al., “DiscDiff: Latent Diffusion Model for DNA Sequence Generation.” Accessed: Jul. 01, 2025. [Online]. Available: http://arxiv.org/abs/2402.06079

[4] A. Fallahpour, V. Gureghian, G. J. Filion, A. B. Lindner, and A. Pandi, “CodonTransformer: a multispecies codon optimizer using context-aware neural networks,” Nature Communications, vol. 16, no. 1, p. 3205, Apr. 2025, doi: 10.1038/s41467-025-58588-7.

[5] “mDD-0: AI Model for mRNA Sequence Generation | Ginkgo Bioworks.” Accessed: Jul. 01, 2025. [Online]. Available: https://ginkgo.bio/resources/white-papers/mrna-discrete-diffusion

[6] Ž. Avsec et al., “Effective gene expression prediction from sequence by integrating long-range interactions,” Nature Methods, vol. 18, no. 10, pp. 1196–1203, Oct. 2021, doi: 10.1038/s41592-021-01252-x.

[7] J. Linder, D. Srivastava, H. Yuan, V. Agarwal, and D. R. Kelley, “Predicting RNA-seq coverage from DNA sequence as a unifying model of gene regulation,” Nature Genetics, vol. 57, no. 4, pp. 949–961, Apr. 2025, doi: 10.1038/s41588-024-02053-6.

[8] Ž. Avsec et al., “AlphaGenome: advancing regulatory variant effect prediction with a unified DNA sequence model.”

[9] D. Penzar et al., “LegNet: a best-in-class deep learning model for short DNA regulatory regions,” Bioinformatics, vol. 39, no. 8, p. btad457, Aug. 2023, doi: 10.1093/bioinformatics/btad457.

[10] J. Zrimec, F. Buric, M. Kokina, V. Garcia, and A. Zelezniak, “Learning the Regulatory Code of Gene Expression,” Frontiers in Molecular Biosciences, vol. 8, Jun. 2021, doi: 10.3389/fmolb.2021.673363.

[11] I. N. Brooks, A. L. Hughes, S. Clauder-Münster, L. A. Mitchell, J. D. Boeke, and L. M. Steinmetz, “Transcriptional neighborhoods regulate transcript isoform lengths and expression levels,” Science, vol. 375, no. 6584, pp. 1000–1005, Mar. 2022, doi: 10.1126/science.abg0162.

[12] I. H. Greger, A. Aranda, and N. Proudfoot, “Balancing transcriptional interference and initiation on the \textit{GAL7} promoter of \textit{Saccharomyces cerevisiae},” Proceedings of the National Academy of Sciences, vol. 97, no. 15, pp. 8415–8420, Jul. 2000, doi: 10.1073/pnas.140217697.

[13] E. M. Prescott and N. J. Proudfoot, “Transcriptional collision between convergent genes in budding yeast,” Proceedings of the National Academy of Sciences of the United States of America, vol. 99, no. 13, pp. 8796–8801, Jun. 2002, doi: 10.1073/pnas.132270899.

[14] E. N. Powers et al., “Bidirectional promoter activity from expression cassettes can drive off-target repression of neighboring gene translation,” eLife, vol. 11, p. e81086, Dec. 2022, doi: 10.7554/eLife.81086.

[15] Z. Jin et al., “Unraveling the regulatory dynamics of bidirectional promoters for modulating gene co-expression and metabolic flux in Saccharomyces cerevisiae,” Nucleic Acids Research, vol. 53, no. 11, p. gkaf511, Jun. 2025, doi: 10.1093/nar/gkaf511.

[16] J. C. Hingerl, A. Karollus, and J. Gagneur, “Flashzoi: An enhanced Borzoi model for accelerated genomic analysis.” Accessed: Jun. 19, 2025. [Online]. Available: https://www.biorxiv.org/content/10.1101/2024.12.18.629121v1

[17] “Transcription Factors and Transcriptional Control | Learn Science at Scitable.” Accessed: Jul. 28, 2025. [Online]. Available: http://www.nature.com/scitable/topicpage/transcription-factors-and-transcriptional-control-in-eukaryotic-1046

[18] Luthra, C. Jensen, X. E. Chen, A. L. Salaudeen, A. M. Rafi, and C. G. de Boer, “Regulatory activity is the default DNA state in eukaryotes,” Nature Structural & Molecular Biology, vol. 31, no. 3, pp. 559–567, Mar. 2024, doi: 10.1038/s41594-024-01235-4.

[19] C. G. de Boer, E. D. Vaishnav, R. Sadeh, E. L. Abeyta, N. Friedman, and A. Regev, “Deciphering eukaryotic gene-regulatory logic with 100 million random promoters,” Nature Biotechnology, vol. 38, no. 1, pp. 56–65, Jan. 2020, doi: 10.1038/s41587-019-0315-8.

[20] A. M. Rafi et al., “A community effort to optimize sequence-based deep learning models of gene regulation,” Nature Biotechnology, pp. 1–11, Oct. 2024, doi: 10.1038/s41587-024-02414-w.

[21] M. E. Lee, W. C. DeLoache, B. Cervantes, and J. E. Dueber, “A Highly Characterized Yeast Toolkit for Modular, Multipart Assembly,” ACS Synthetic Biology, vol. 4, no. 9, pp. 975–986, Sep. 2015, doi: 10.1021/sb500366v.

[22] O. Shalem et al., “Systematic Dissection of the Sequence Determinants of Gene 3’ End Mediated Expression Control,” PLOS Genetics, vol. 11, no. 4, p. e1005147, Apr. 2015, doi: 10.1371/journal.pgen.1005147.

[23] C. Lu, L. Guo, B. Fang, J. Shi, and M. Zhou, “DNA Sequence Changes Resulting from Codon Optimization Affect Gene Expression in Pichia pastoris by Altering Chromatin Accessibility,” ℱournal of Fungi (Basel, Switzerland), vol. 11, no. 4, p. 282, Apr. 2025, doi: 10.3390/jof11040282.

[24] Schreiber, F. K. Lorbeer, M. Heinzl, Y. Y. Lu, A. Stark, and W. S. Noble, “Programmatic design and editing of cis-regulatory elements.” Accessed: Jul. 25, 2025. [Online]. Available: https://www.biorxiv.org/content/10.1101/2025.04.22.650035v1

[25] Su, Y. Lu, S. Pan, A. Murtadha, B. Wen, and Y. Liu, “RoFormer: Enhanced Transformer with Rotary Position Embedding.” Accessed: Jun. 29, 2025. [Online]. Available: http://arxiv.org/abs/2104.09864

[26] D. Takai and P. A. Jones, “Origins of bidirectional promoters: computational analyses of intergenic distance in the human genome,” Molecular Biology and Evolution, vol. 21, no. 3, pp. 463–467, Mar. 2004, doi: 10.1093/molbev/msh040.

[27] Ž. Avsec et al., “Base-resolution models of transcription-factor binding reveal soft motif syntax,” Nature Genetics, vol. 53, no. 3, pp. 354–366, Mar. 2021, doi: 10.1038/s41588-021-00782-6.

[28] A. Karollus, J. Hingerl, D. Gankin, M. Grosshauser, K. Klemon, and J. Gagneur, “Species-aware DNA language models capture regulatory elements and their evolution,” Genome Biology, vol. 25, no. 1, p. 83, Apr. 2024, doi: 10.1186/s13059-024-03221-x.

