## Supplementary Materials for "Yorzoi: Predicting RNA-seq coverage from DNA sequence in yeast"

### Supplementary Material

#### Figure S1: Bias introduced by sample splitting

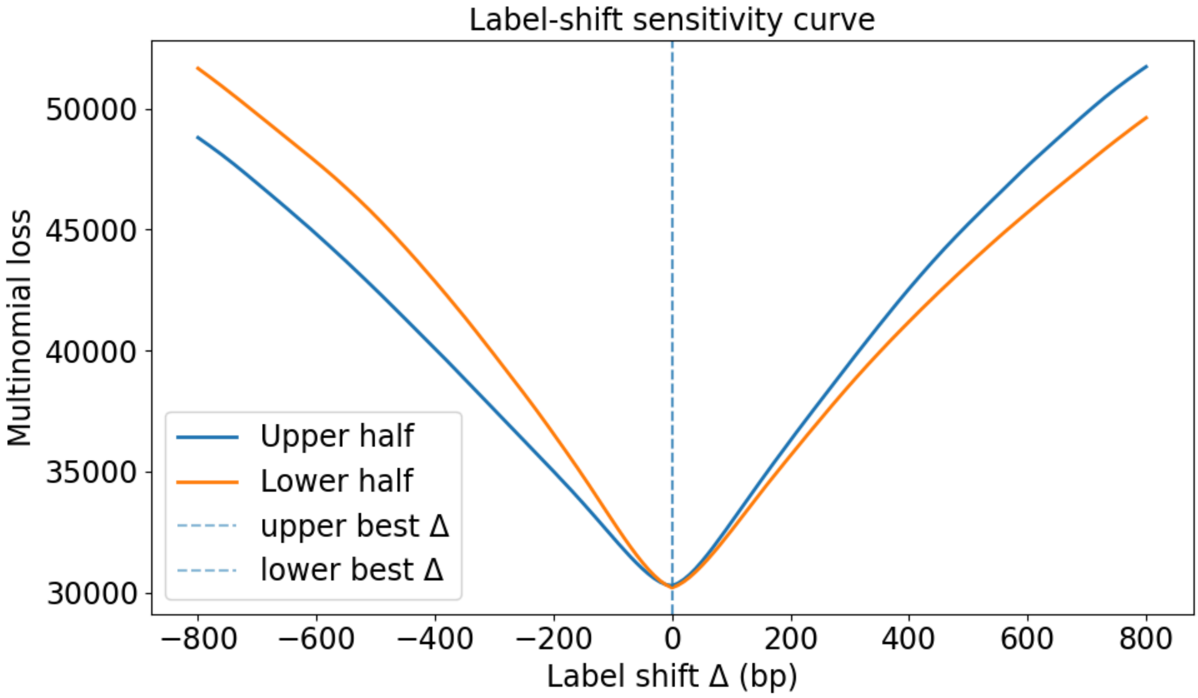


Figure 1:

**Position-bias check.** Our 3 kb windows (±1 kb context) may induce an absolute-position prior. We test this with a label-shift sensitivity analysis. Model the normalized prediction as

$$p_{\text{pred}}(k) \approx (1-\alpha)q(k \mid x) + \alpha b(k)$$

Keeping $p_{\text{pred}}$ fixed, we circularly shift the labels by $\Delta$ bins within the 3 kb window and recompute the multinomial (shape) loss:

$$L(\Delta) = -\sum_k y^{\Delta}(k) \cdot \log p_{\text{pred}}(k)$$

with

$$y^{\Delta}(k) = y(k-\Delta)$$

A position template would move the minimum to $\Delta \neq 0$; in our data both halves minimize at $\Delta = 0$ indicating negligible position-offset bias.

### DREAM Promoter Sequence Construction

```
import pandas as pd
import torch
from torch.utils.data import Dataset
from yorzoi.dataset import GenomicDataset


class PromoterDataset(Dataset):
    upstream_polyT_777bp =
"gcggcagaagaagtaacaaaggaacctagaggcctttttgatgttagcagaattgtcatgcaagggctccctatctactggagaatatactaagggtac
    polyT_no_overhang = "gctagcaggaatgatgcaaaggttcccgattcgaac"
```

```
    polyA_no_overhang = (

"tcttaattaaaaaaagatagaaaacattaggagtgtaacacaagactttcggatcctgagcaggcaagataaacga"
    )
    YFP_connector = "aggcaaag"
    YFP =
"atgtctaaaggtgaagaattattcactggtgttgtcccaattttggttgaattagatggtgatgttaatggtcacaaattttctgtctccggtgaagg
    YFP_term_connector = "ggcgcgccacttctaaataa"
    ADH1_terminator =
"gcgaatttcttatgatttatgatttttattattaaataagttataaaaaaaataagtgtatacaaattttaaagtgactcttaggttttaaaacgaaa
    term_natmx6_connector = (
        "cagatccgctagggataacagggtaatatagatctgtttagcttgcctcgtccccgccgggtcacccggccagc"
    )
    natmx6 =
"gacatggaggcccagaataccctccttgacagtcttgacgtgcgcagctcaggggcatgatgtgactgtcgcccgtacatttagcccatacatcccca
    natmx6_cen_connector = (
        "ctgtcgattcgatactaacgccgccatccagtgtcgaaaacgagctcgaattcctgggtccttttc"
    )
    cen =
"atcacgtgctataaaaataattataatttaaatttttaatataaatatataaattaaaaatagaaagtaaaaaaagaaattaaagaaaaaatagttt
    cen_downstream =
"cgtccgatatcatcagatccactagtggcctatgcggccgcggatctgccggtctccctatagtgagtcgtattaatttcgataagccaggttaacct

    polyT_overhang = "TGCATTTTTTTCACATC".lower()
    polyA_overhang = "GGTTACGGCTGTT".lower()

    def __init__(self, promoter_seq_file: str):
        self.data = pd.read_csv(
            promoter_seq_file, sep="\t", header=None, names=["seq", "el"]
        )

    def construct_sequence(self, n80_with_overhang: str):
        n80_with_overhang = n80_with_overhang.lower()

        assert (
            len(n80_with_overhang) == 110
        ), f"n80 with overhang is not 110bp long. {len(n80_with_overhang)}"

        assert n80_with_overhang.startswith(
            PromoterDataset.polyT_overhang
        ), f"Seq doesn't start with polyT overhang! PolyT:
{PromoterDataset.polyT_overhang} but got {n80_with_overhang[:
len(PromoterDataset.polyT_overhang)]}"
        assert n80_with_overhang.endswith(
            PromoterDataset.polyA_overhang
        ), f"Seq doesn't start with polyT overhang! PolyA:
{PromoterDataset.polyA_overhang} but got {n80_with_overhang[-
len(PromoterDataset.polyA_overhang) :]}"

        full_seq = (
            PromoterDataset.upstream_polyT_777bp  # 777bp
            + PromoterDataset.polyT_no_overhang  # 37 bp
            + n80_with_overhang  # 110 bp
            + PromoterDataset.polyA_no_overhang  # 76 bp
            + PromoterDataset.YFP_connector  # 8 bp
            + PromoterDataset.YFP  # 717 bp, starts at 1008 bp
```

```
            + PromoterDataset.YFP_term_connector
            + PromoterDataset.ADH1_terminator
            + PromoterDataset.term_natmx6_connector
            + PromoterDataset.natmx6
            + PromoterDataset.natmx6_cen_connector
            + PromoterDataset.cen
            + PromoterDataset.cen_downstream
        )

        assert (
            len(full_seq) == 5000
        ), f"Sequence is not 5 Kb long. Seq len {len(full_seq)}"

        return full_seq

    def __len__(self):
        return len(self.data)

    def __getitem__(self, idx):
        construct_sequence = self.construct_sequence(self.data.iloc[idx]
["seq"]).upper()

        return (
            torch.tensor(
                GenomicDataset.one_hot_encode(construct_sequence),
dtype=torch.float32
            ),
            torch.tensor([self.data.iloc[idx]["el"]]),
        )
```
